# Non-covalent Linking Antibody to Yeast Using the Cell Surface Display of Staphylococcal Protein A

**DOI:** 10.1101/2023.06.30.547162

**Authors:** Yi-Feng Shi

## Abstract

Yeast surface display represents a commonly used platform suitable for the generation and screening of antibodies as well as the selection of high-producer clones. The methods of yeast display rely on the genetic fusion of recombinant antibodies to an abundant cell wall protein of yeast. Here, the study of proof of concept showed that the conventional strategy of expression of fusion antibodies was replaced by non-covalent binding to antibodies for yeast surface display. The use of cell surface display of an epitope tag will endow the cells with new arms for immobilizing, absorbing or targeting the proteins. *Staphylococcal* protein A, which is characterized by its ability to bind selectively to the Fc region of IgG, was examined to be expressed on the surface of *Saccharomyces cerevisiae* using a secretion signal of *Rhizopus oryzae* glucoamylase and C-terminal half of α−agglutinin including glycosylphosphatidylinositol (GPI) anchor attachment signal under the control of the glyceraldehyde 3-phosphate dehydrogenase (*GAPDH*) promoter. On the other hand, an Fc-fused enzyme was created to construct a molecular fusion of *Rhizopus oryzae* Lipase with the spacer and the Fc region of IgG heavy chain. The secretion of fusion protein was carried out using pre-α-factor leader region as secretion signal under the control of the 5^’^-upstream region of the *Candida tropicalis* isocitrate lyase gene (*UPR-ICL*) in *S. cerevisiae*. The Fc-fused lipase was captured by Staphylococcal protein A as an adaptor protein displayed on the surface of yeast cells. The method of this switchable yeast display takes advantage of the “secretion-and-capture” strategy and can be applied to improve the efficiency of yeast display of full-length IgG.

**graphical abstract:** Staphylococcal protein A, which has the ability to binding to the Fc region of IgG and leaving the antigen combing site free, has been assembled on the surface of yeast cells to target antibodies or enzymes with Fc fusion.

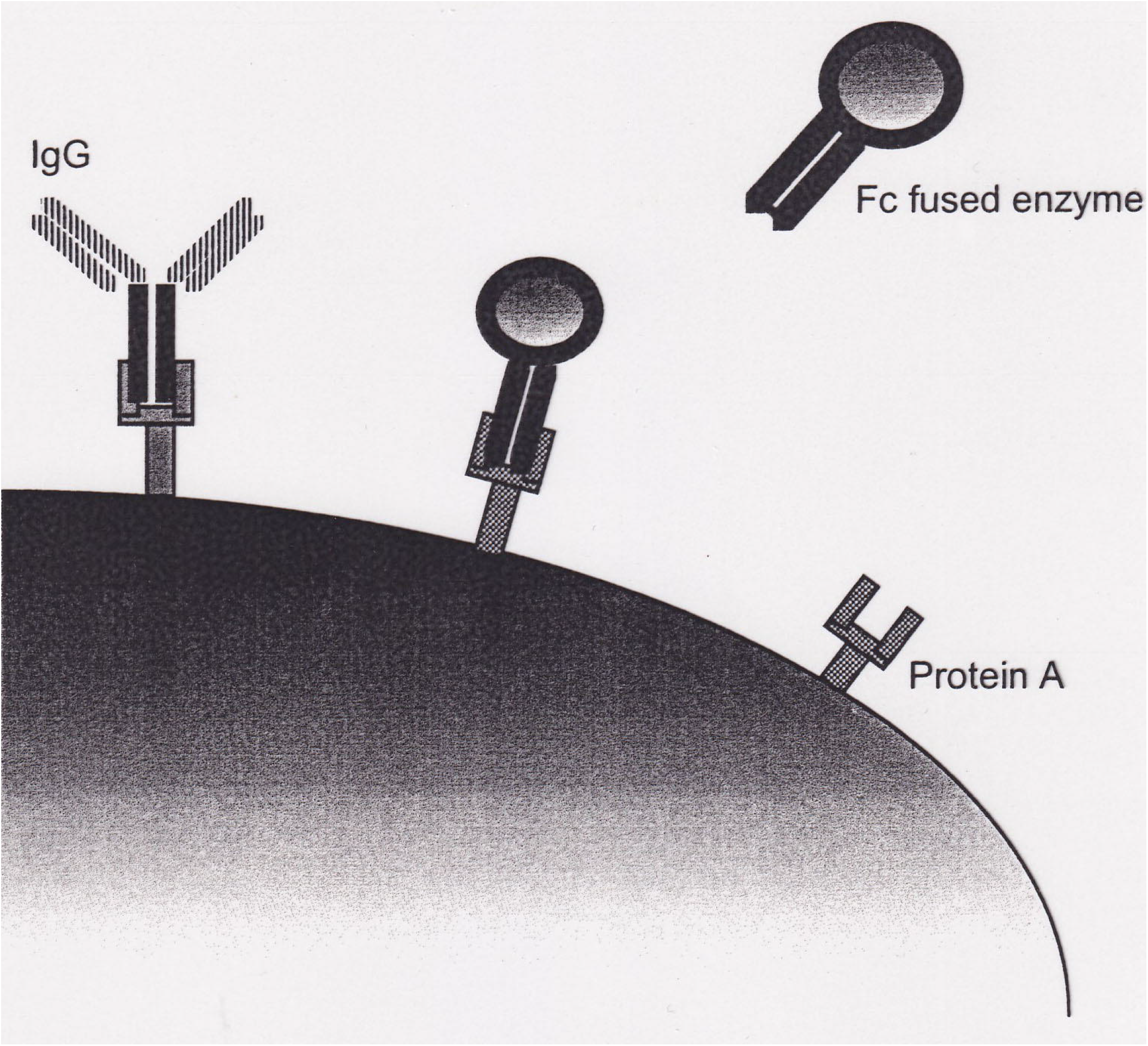

## Introduction

Antibodies are ideal therapeutic and diagnostic molecules, due to their very high specificity and selectivity against targets of interest (*Chiu & Gilliland,2016*). Although animal immunization was initially employed as an antibody discovery approach, combinatorial biology methods, such as phage or yeast display technology, which is based on in vitro selection from molecular libraries and overcoming limitations of immune tolerance, have transformed the development of monoclonal antibodies. Using yeast as eukaryotic cell expression can be expected to display better properties of well-folded and glycosylated antibodies compared to bacteria protein production for phage display (*Cherf & Cochran, 2015; Könning & Kolmar, 2018; Teymennet-Ramírez, 2022*).

Staphylococcal protein A is a cell-wall-bound pathogenicity factor from *Staphylococcal aureus*. It exhibits tight binding to the Fc region of immunoglobulins IgG, leaving the antigen combing sites free. This ability has long been utilized as a powerful immunological tool (*Mouratou et al. 2015; Bolton & Mehta,2016; von Witting et al. 2021*). The gene for protein A encodes a total of 254 residues, including an N-terminal signal sequence, five highly homologous~58 residue IgG binding domains, and a C-terminal region containing the site for cell wall attachment (*Uhlen, et al*. *1984*; *Moks, et al. 1986*). Because of their small size and IgG-binding activity, domains of protein A are targets for protein engineering. The Z domain, a more stable engineered analog of the B domain, is an antiparallel three-helix, 59-residue bundle that has been studied and developed deeply for understanding determinants of protein function and stability (*Nilsson et al. 1987; Braisted & Wells 1996; Nord et al, 1997*). For the purpose of screening novel binding proteins from combinatorial libraries and development of live vaccine vehicles, many expression systems for surface display of staphylococcal protein A have been studied. Chaudhary and coworkers have reported phage display of functional Ig-G binding domains of staphylococcal protein A (*Chwaha et al*., *1994*). Meruelo and colleagues constructed a Sinbis virus vector displaying the IgG-binding domain (*Ohno et al*., *1997*). Remaut and coworkers showed that the cell wall anchor of protein A from *Staphylococcal aureus* is functional in the *Lactococcus lactis* for display of a heterologous protein (*Steidler et al*., *1998*). Kronqvist and colleagues have displayed the library of staphylococcal protein A (SpA)-derived affibody molecules on the surface of staphylococci and used the flow cytometry to isolate binders with subnanomolar apparent affinity for TNF-alpha successfully (Kronqvist et al., 2008). Vahed and colleagues experimental studied the surface display of SPA protein via Lpp-OmpA system for the screening of IgG (Vahed et al., 2020).

Yeast surface display of heterologous proteins has been utilized for various applications which include recombinant vaccines, polypeptide libraries, whole-cell adsorbents and recombinant whole-cell biocatalysts (*Kondo & Ueda, 2004;Shibasaki & Ueda 2014; Kuroda & Ueda, 2022;Murai et al. 1999; Schreuder et al. 1996; Georgiou et al. 1997; Stahl & Uhlen, 1997*). Lipases (EC 3.1.1.3) are capable of hydrolyzing the ester bonds of hydrophobic substrates. They have wide specificity of substrate and can also catalyze ester synthesis in reverse direction. They have been widely used in food, pharmaceutical and detergent industries (*Jaeger & Reetz, 1998*; *Schmid & Verger*,. *1998*). Washida, Matsumoto, et al. have constructed a *Saccharomyces cerevisiae* strain displaying an active lipase on the cell surface by GPI anchor and the Flo1p flocculation functional domain respectively (*Washida et al. 2001: Matsumoto et al. 2002*)

Epitope tagging makes the recombination protein immunoreactive to a known antibody by fusing epitope (*Jarvik & Telmer, 1998*). Its important feature is that it provides a specific means to separate, locate and target the tagged proteins or tagged cells. Staphylococcal protein A as an epitope tag has been widely exploited for purification or immobilization of recombinant protein on IgG affinity solid support (*Nilsson et al*., *1985*; *Nilsson et al*., *1994*). Utilizing such strong and highly specific interaction between Staphylococcal protein A and Fc region of IgG, we can design a novel enzyme immobilization method combined with cell surface display technology.

Here, two vectors for engineering genes of yeast display of staphylococcal protein A on the cell surface and a molecular fusion of lipase with the Fc region of IgG were constructed respectively. The creating lipase-Fc fusion enzyme can be targeted by staphylococcal protein A on the surface of yeast cells. The method of this switchable yeast display can be developed universally to improve the efficiency of yeast display of full-length IgG.

## Materials & Methods

### Strains and media

*Escherichia coli* DH5α [F^−^ *end*A1 *hsd*R17 (r_K_^−^ m_K_^−^) *supE44 thi-1* λ^−^ *recA1 gyrA96 ΔlacU96*(Φ80*lac*Z-ΔM15) was used as a host for recombinant DNA manipulation. *Saccharomyces cerevisiae* MT8-1 (*MAT***a** *ade hie3 leu2 trp1 ura3*) was used for expression of staphylococcal protein A and Fc-fused lipase. Transformants of *E. coli* were grown in LB medium (1 % tryptone, 0.5 % yeast-extract, 0.5 % sodium chloride) containing 0.1 % glucose and 50 μg/ml ampicillin at 37 °C. The transformants of yeast were selected on the SD medium (0.67 % yeast nitrogen base without L-Tryptophan, 2% glucose and appropriate supplement) at 30 °C. The transformants of yeast were cultured in YPD medium (1 % yeast extract, 2 % peptone, 2 % glucose) and further cultured in YAP medium (1 % yeast estract, 2 % peptone, and 1 % sodium acetate.

### Construction of the plasmids for cell surface display of staphylococcal protein A

The plasmid pRIT2T was used as the donor of the gene coding for Staphylococcal protein A (*Nilsson et al*., *1985*). The plasmid pICAS1 and pCAS1 were used as cassette vectors (*Murai et al*., *1998*) for the expression of staphylococcal protein A on cell surface. The primers (5’-AAGAAG<u>CCGCGG</u>ATCAACGCAATGGTTTT-3’ and 5’-TTTATC<u>CTCGAG</u>TTTGTTATCTGCAGGTCGACGG-3’) were designed to introduce *Sac*II and *Xho*I cloning sites in the ends of the gene fragment of staphylococcal protein A respectively in PCR amplification with pRIT2T as the template. Other genes except for the protein A-encoding gene were truncated and stop code was added in the staphylococcal protein A-encoding gene fragment. The gene of staphylococcal protein A was then cloned into pICAS1 at their *Sac*II and *Xho*I cloning sites to construct pICASA fusion vectors which were checked by DNA sequence analysis.

### Construction of the plasmids for Fc-fused Lipase

The PCR method was used to introduce *Bgl* II and *Sal* I cloning sites at the 5’-end and *Xho*I cloning site at 3’-end of the Fc portion of human IgG by the Primers of 5’-ACTCAC<u>AGATCT</u>CCACCG<u>GTCGAC</u>GCACCTGAACTCCTGG-3’and 5’-CTACTA<u>CTCGAG</u>TCATTTACCCGGAGACAAGGGAGAGGC-3’ respectively. Such Fc fragment of human IgG was cloned into the cassette vector pWI3 at the cloning sites of *Bgl*II and *Xho*I to construct the plasmid pWI-Fc. The *Rizopus oryzae* lipase gene (*Takahashi et al*.,*1998*) having the peptide linker (GSSGGSGGS) was introduced *Bgl*II and *Sal*I cloning site by PCR method using the primers of 5’-AGTTTC<u>AGATCT</u> ATGAGATTTCCTTCAATTTTTACTGC-3’ and 5’-GCTTTT<u>GCTGAC</u>CGAACC ACCAGAACCACC-3’and was then cloned into the plasmid of pWI-Fc at the *Bgl*II and *Sal*I cloning site. The resultant plasmid named pWI-Fc-ROL was checked by DNA sequence analysis.

### Expression of recombinant proteins

The transformation of *S. cerevisiae* was carried out by using the lithium acetate method of Yeastmaker™ Yeast Tansformation System (CLONTECH Laboratories, Inc.). The plasmid of pICASA was cleaved by *Xba*I and integrated into the chromosomal DNA of *S. cerevisiae* and selected on the SD-Trp medium. The circular plasmids and pWI-Fc-ROL were transformed into *S. cerevisiae* selected on SD-Trp medium. The transformation yeasts harboring pICASA were cultured aerobically in YPD medium for 1 day at 30°C to express Staphylococcal protein A on the cell surface. The transformation yeasts of pWI-Fc-ROL were precultured aerobically in YPD medium and further cultivated in YAP medium at 30°C for 2 day to express and secrete Fc-fused lipase.

### Immunofloresence microscopy

For immunostaining, washed, overnight cells were resuspended in phosphate buffer saline (PBS) containing 1% bovine serum albumin and incubated for half an hour. Subsequently the cells were pelleted by centrifugation, washed and then incubated in the same PBS/BSA solution with mouse protein A antibody at 1:1000 dilution for 1 hour. Following another two washes with PBS the cells and exposed to the second antibody, fluoresein-isothiocyanate-conjugated goat anti-mouse IgG, diluted 1:300 at room temperature for 1 hour. Finally, the cells were resuspended in PBS and examined by fluorescence microscopy.

### SDS-PAGE and Western Blotting

To evaluate the production of recombination fused proteins, using late exponential phase cells (OD_600_ 1.0) was analyzed by SDS-PAGE on 12 % acrylamide gels and by Western blotting, anti-protein A or anti-lipase antibodies, horseradish peroxidase-conjugated the second antibodies and the peroxidase substrate 4-chloro-1-paphtol stationing solution.

#### Enzyme assay

Plate assay was performed on YAP agar plate containing 1 % tributyrin (Wako, Osaka) emulsified by sonication and 25 μg/ml ampicillin. Lipase activity was measured with Lipase Kit S (Dainippon Pharmaceutical, Osaka, Japan) at 30°C. One unit of the enzyme activity was defined as the amount of the enzyme that catalyzes the formation of 1 μmol of 2,3-dimercaptopropan-1-ol from 2,3-dimercaptopropan-ol tributyl ester per min.

## Results

### Staphylococcal protein A to express on the surface of *Saccharomyces cerevisiae* cell

The integrative plasmid of pICASA was constructed for the display Staphylococcal protein A on the surface of *Saccharomyces cerevisiae* (Figure 1). in which we used a secretion signal of *Rhizopus oryzae* glucoamylase and C-terminal half of α−agglutinin including glycosylphosphatidylinositol (GPI) anchor attachment signal under the control of the glyceraldehyde 3-phosphate dehydrogenase (*GAPDH*) promoter. The cell wall mannoprotein α−agglutinin, which is involved in the sexual adhesion of mating-type *S. cerevisiae* and mating-type a *S. cerevisiae*, is covalently linked to cell wall glucan which can only be split by the enzymes like glucanase but not by SDS extraction. The anchoring ability is conferred by the C-terminal half of protein which has a hydrophobic tail but replace by GPI anchor in the endoplasmic reticulum.

**Figure 1.**
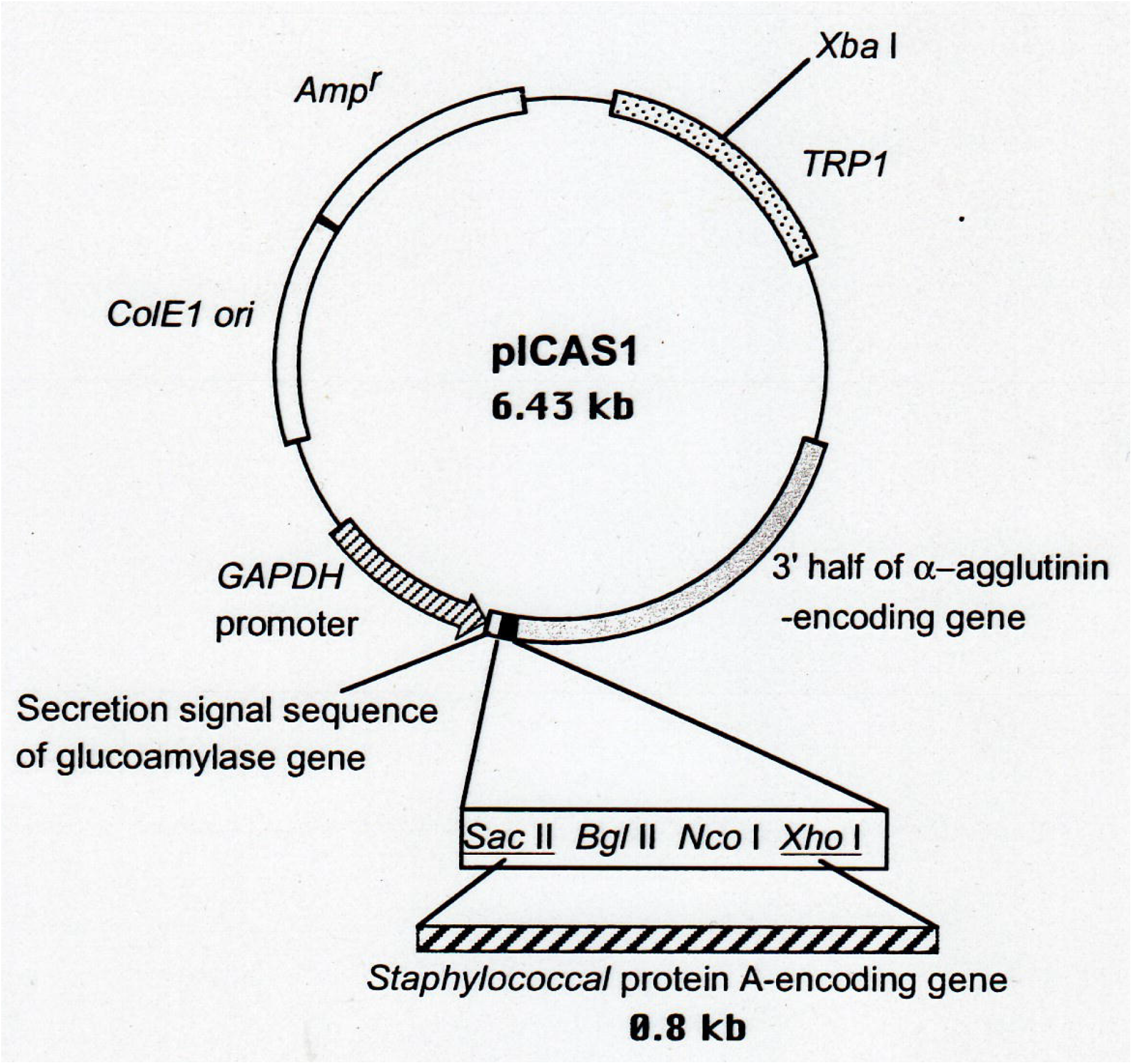
Integrative plasmid, pICASA, to display Staphylococcal protein A on the surface of *Saccharomyces cerevisiae*.

For the tansformation of the plasmid pICASA into *S. cerevisiae*, integration can occur at *TRP1* locus by homologous recombination. *TRP1* gene of pICASA was cut by *Xba*I and then the transformation of *S. cerevisiae* was carried out using the lithium acetate method. The synthetic medium with tryptophan omitted was used to select the positive transformants. Integration of pICASA encoding staphylococcal protein A into the chromosome of *S. cerevisiae* was verified by PCR method in which the staphylococcal protein A gene was amplified using the genome DNA of *S. cerevisiae* transformant as the template (Figure 2).

**Figure 2.**
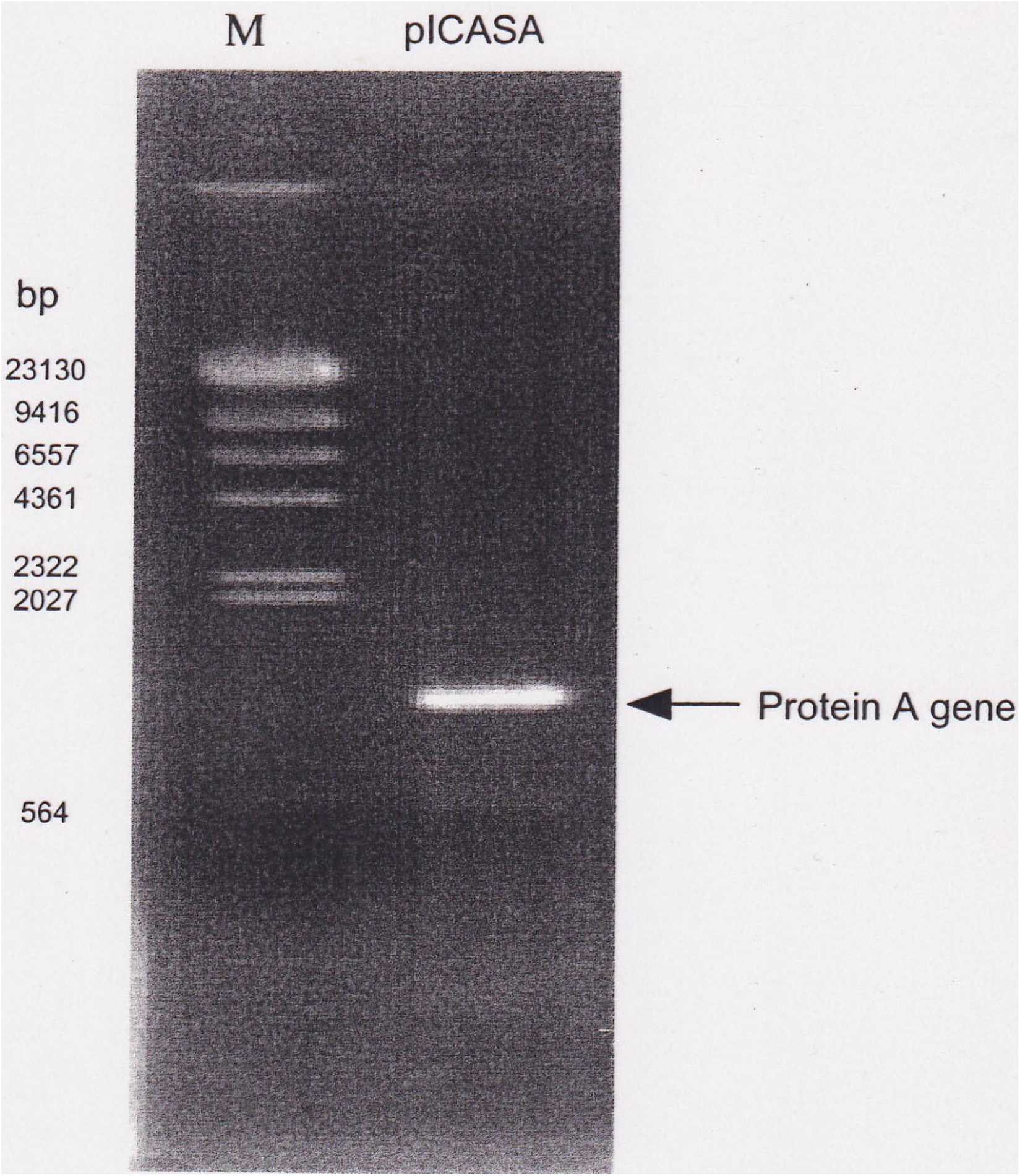
Demonstration of the integrated Staphylococcal protein A-encoding gene by PCR.

### *Rhizopus oryzae* Lipase to fuse to the Fc region of IgG

We constructed a molecular fusion of *Rhizopus oryzae* Lipase with the spacer and the Fc region of the human IgG heavy chain, creating an Fc-fused enzyme (Figure 3). The plasmid pWI3 was used as the cassette vector. The lipase gene and Fc region of human IgG were cloned into it to construct pWI-lipase-Fc (pWI-ROL-Fc) plasmid shown in Figure 3 for expression and secretion of Fc-fused lipase. Lipase produced by *Rhizopus Oryzae* (ROL) comprise a signal sequence of 26 amino acids, a prosequence of 97 amino acids, and a mature lipase region of 269 amino acids (*Beer et al. 1996*). We use the 5^’^-upstream region of the *Candida tropicalis* isocitrate lyase gene (*UPR-ICL*) as a promoter for high-level expression of lipase. The pre-a-factor leader region of *S. cerevisiae* replaced the signal sequence of ROL to assist the secretion of the lipase. The carboxyl terminus of lipase was fused to the Fc domain (C_H_2 and C_H_3) of human IgG1 due to the characterization of staphylococcal protein A binding region. The spacer of nine peptides was placed between Fc and lipase in taking consideration of their structure folding or integration. The plasmid of pWI-ROL-Fc was checked by digestion of *Xho*I, *Bgl*II and *Sal*I shown in the electrophoresis (Figure 4.)

**Figure 3.**
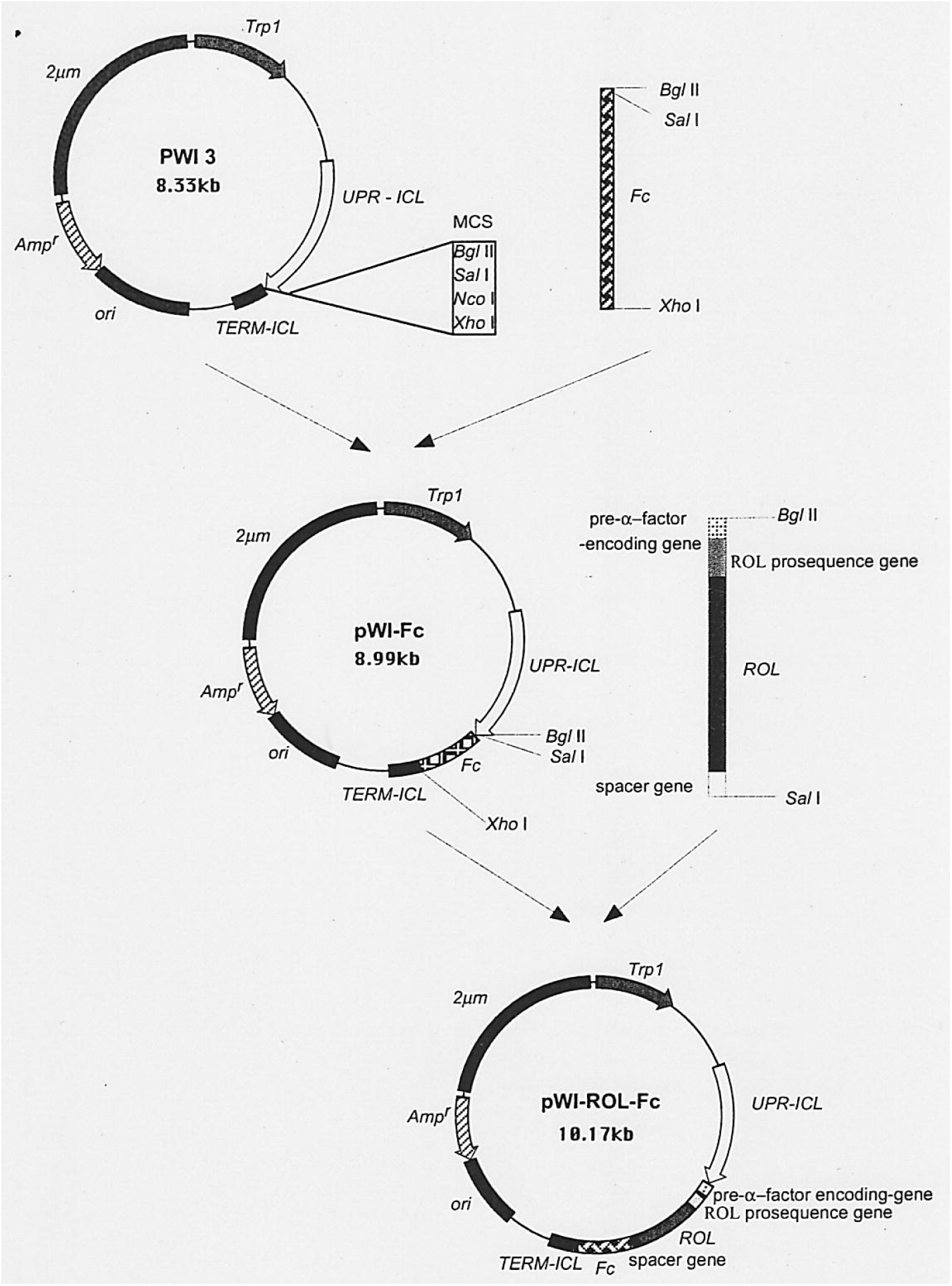
Construction of pWI-ROL-Fc to express the Fc-fused lipase.

**Figure 4.**
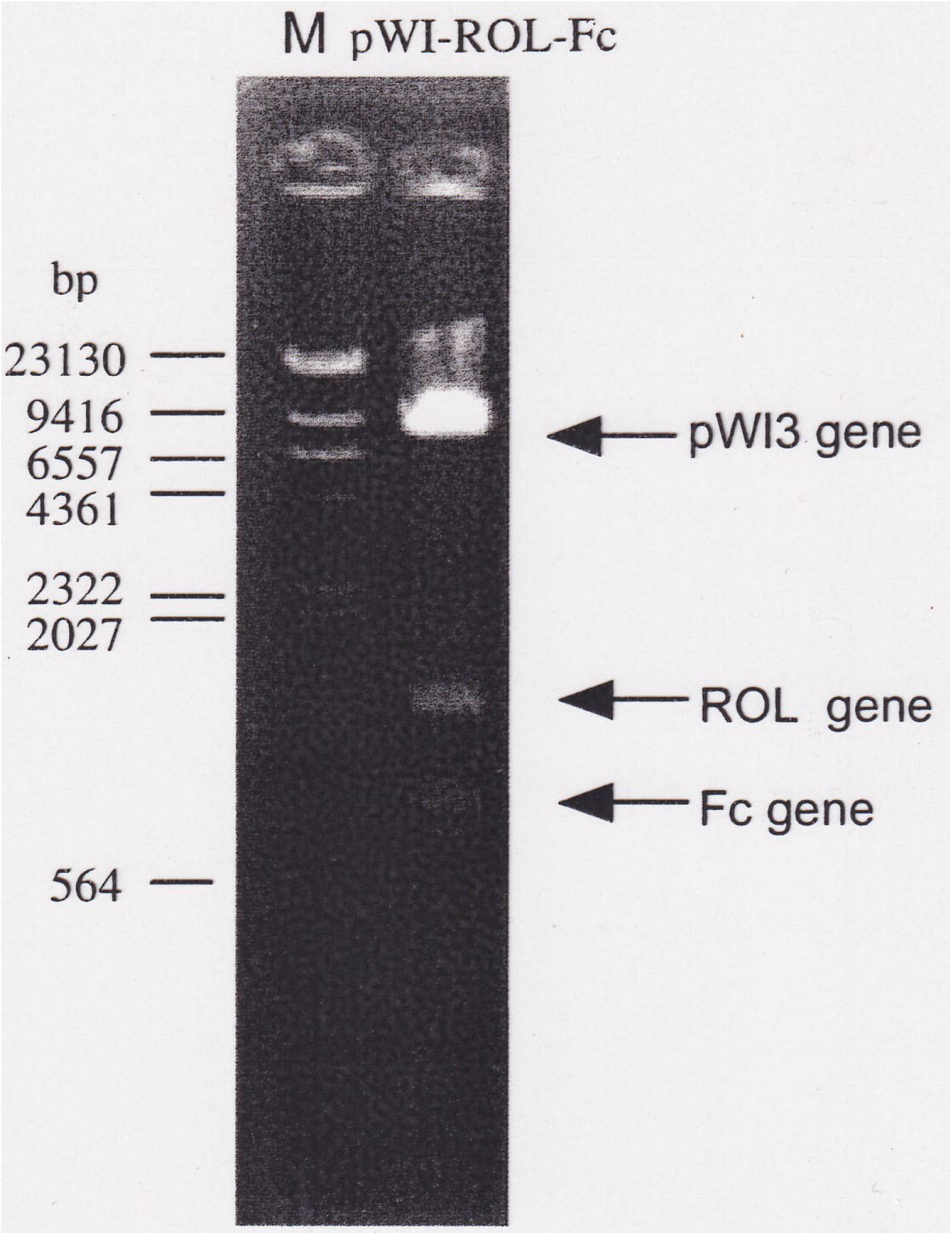
The plasmid of pWI-ROL-Fc was checked by digestion of *Xho*I, *Bgl*II and *Sal*I shown in the electrophoresis

### Evaluation of expression of the constructed genes

Immunofluorescence labeling was applied to identify the production and localization of protein A expression on the surface of *S. cerevisiae*. Using an anti-protein A antibody as the first antibody and anti-IgG antibody with FITC as the second antibody, the expression of protein A on the cell surface can be examined by immunofluorescence microscopy.

We performed a plate assay to detect the transformants with the lipase activity. The results in Figure 5 demonstrated that the transformants hydrolyzed the substrate tributyrin and formed holo around clones. The transformation *S. cerevisiae* harboring pWI-Fc-ROL were precultured aerobically in YPD medium and further cultivated in YAP medium at 30°C for 2 day to express and secrete Fc-fused lipase. The lipase activity in the culture supernatants was measured. Western blotting was applied to analysis Fc-fused lipase and its binding activity with protein.

**Figure 5.**
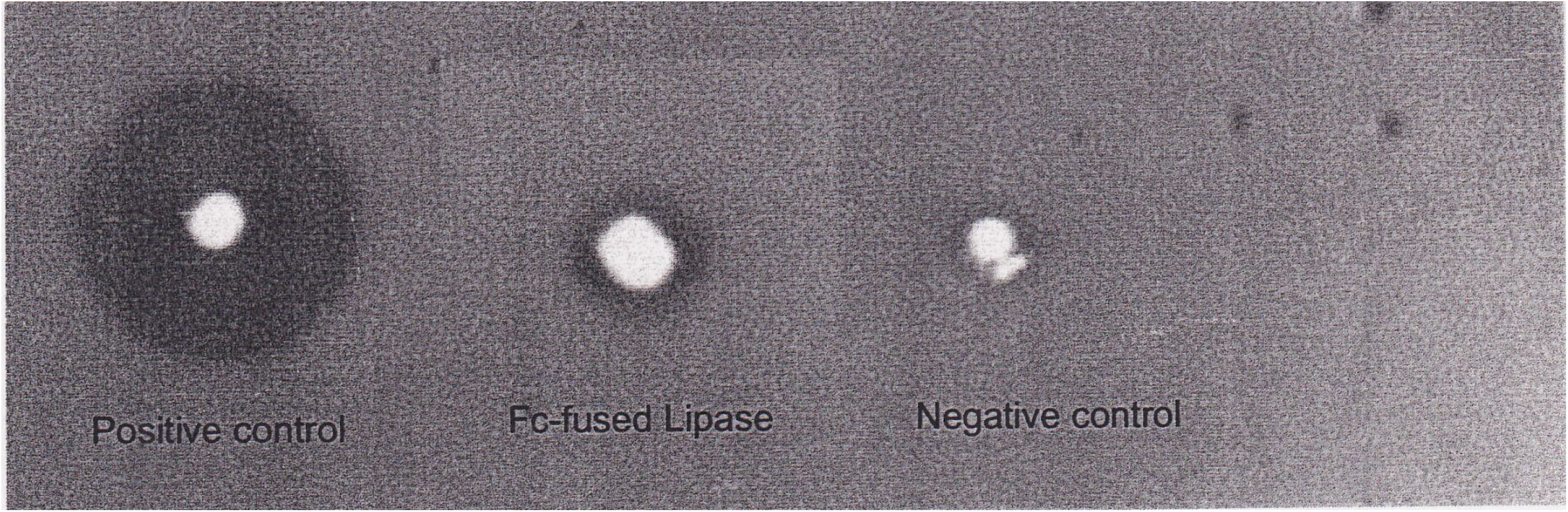
A plate assay to detect the transformants with the lipase activity. (Holo around clones from left to right: Positive control; Fc-fused lipase; negative control)

## Discussion

Due to the presence of several cell-wall anchored wall proteins, the selection of the anchor protein became a crucial genetic strategy. The most common cell wall-anchoring proteins are α−agglutinin and a-agglutinin which are two mating specific agglutinin that mediate cell-cell contact of α and a mating types. During formation of the cell wall, the fusion protein is targeted to the cell surface through its C-terminal GPI (glycosylphosphatidylinositol)-attachment signal. The displayed proteins are often fused to the secretion signal sequence at N-terminus and to the half α−agglutinin domain at the C-terminus. On the other hand, a-agglutinin-based anchor, which is Aga1-Aga2 linked by disulfide bond, allow the immobilization of proteins through their N- and C-terminal In addition, Flo1p, which is a GPI-anchored cell wall protein involved in flocculation, also provides N-terminal-free molecular display. Recently the Flo1-derived anchor protein has been successfully used to yeast surface display of ZZ domain of staphylococcal protein A (*Katsurada et al*., *2021*).

The new format of yeast display described here is unique in that it takes advantage of binding of staphylococcal protein A to the Fc domain of antibody. Based on the “secretion-and-capture” strategy, the genes of the Fc domain and staphylococcal protein A were genetically fused to two vectors for protein expression of soluble form and cell-surface displayed form respectively. Shibasaki and colleagues reported that the ZZ-domain-displaying cell and Fc fusion lipase-secreting cell can be applied to use in synergistic process of production and recovery of secreted recombinant proteins (*Shibasaki et al. 2012*). Despite a potential loss of genotype-phenotype linkage if using two yeast hosts, there is the major advantage that it is possible to switch between antibody display for screening and antibody secretion for functional characterization of individual candidate antibodies. Furthermore, such staphylococcal protein A mediated binding of antibody to yeast may be coupled with flow cytometric analysis to screen very high diluted antibodies. Because cells are large enough to be screened using high-speed flow cytometers, with the high polyvalency on the cell surface, it enables a real-time quantitative screening of the affinity of all individual library members. The affinity of the candidate clones can rapidly be determined on-cell using flow cytometry without any need for laborious subcloning and protein purification.

## Acknowledgment

The author thanks Prof. Atsuo Tanaka and Mitsuyoshi Ueda of Kyoto University for their suggestion and support on yeast cell surface engineering.

